# Manipulation of encapsulated artificial phospholipid membranes using sub-micellar lysolipid concentrations

**DOI:** 10.1101/2023.06.25.546396

**Authors:** Pantelitsa Dimitriou, Jin Li, William D. Jamieson, Johannes J. Schneider, Oliver K. Castell, David A. Barrow

## Abstract

Droplet Interface Bilayers (DIBs) constitute a commonly used model of artificial membranes for molecular biology studies with applications in synthetic biology research. However, these model membranes have limited accessibility due to their requirement to be surrounded by an oil environment. Here, we demonstrate in-situ bilayer manipulation of submillimeter, free-standing, encapsulated droplet interface bilayers (eDIBs) in hydrogel capsules formed using dual-material, 3D-printed microfluidic devices. These microfluidic devices required no post-fabrication assembly, nor surface treatment to achieve the high-order emulsification, required for the formation of robust eDIBs. The eDIB capsules were exposed to various concentrations of lysophosphatidylcholine (LPC), in order to investigate the interaction of lysolipids with three-dimensional, encapsulated droplet bilayer networks. Micellar LPC concentrations trigger the bursting of the eDIB droplets, while at concentrations below the critical micelle concentration (CMC), the encapsulated aqueous inner droplet networks endure structural changes, precisely affecting the DIB contact angles and bilayer area. Manipulation of these enclosed, 3D-orchestrated membrane mimics facilitates the exploration of readily accessible compartmentalized artificial cellular machinery. Collectively, the multi-compartmentalized capsules and the lysolipid-mediated membrane modulation, introduce a chemical approach to control the properties and mechanics of artificial cellular membranes, toward responsive soft material developments.

## Introduction

Droplet interface bilayers (DIBs) are bottom-up, cellular membrane-mimicking models used for the in-vitro study of membrane constituents and properties ^1^. DIBs are formed when lipid monolayer-coated aqueous droplets come into contact, forming an artificial lipid bilayer membrane. In addition, DIBs can be formed when an aqueous droplet sits on top of a hydrogel substrate ^2,3^, where this model has been used in single-molecule imaging for biophysical and biochemical studies ^4,5^. The versatility of DIB models enables them to be tailored for different research applications, ranging from the study of transmembrane protein behaviour ^2^, to cell-free DNA expression ^6^ and in-vitro tissue culture development ^7^.

Sophisticated and functional artificial cellular networks can be constructed using DIBs as building blocks. Multisomes ^8^, enclose DIB networks within an oil droplet, which can be suspended in air or water ^9–11^. Various multisome demonstrations have been assembled using liquids only ^8,11^, although, the encapsulation of DIBs and multisomes within soft hydrogels ^12^, introduces soft material platforms towards the study of artificial membranes. Hydrogels are attractive because they are used for the immobilization of biological and non-biological matter, including living cells and synthetic cells, respectively ^13,14^.

DIBs on hydrogel substrates acquire enhanced mechanical resistance leading to their prolonged stability and extended lifetime ^12,15,16^. Gel-encapsulated droplet interface bilayer constructs (eDIBs) ^17,18^ are a type of multisomes, which depict multi-compartmentalized artificial cell chassis and aim to impart cellular functionalities, such as polarization ^19^. Furthermore, DIB systems are usually made by manual pipetting ^20^, which limits the production yield rate and structural complexity attained. Recently, multiphase microfluidic droplet-forming devices have been developed to effectively generate DIBs, multisomes and eDIBs, using stepwise emulsification methods ^8,17,19^. Such droplet-based artificial membrane networks formed by robust and high-throughput microfluidic techniques have been used in molecular sensing ^8^, cell mimicking ^19^, and artificial cell membrane studies ^17^. The properties of simple and complex DIB systems are determined largely by the lipid and oil composition ^21^, membrane chemistry ^22^, as well as the droplet arrangement ^19,23^. Bilayer mechanics, forces and capacitance are characteristics directly influenced by the conditions of a DIB model ^24–26^. Various studies have focused on the geometrical parameters of DIBs, e.g., contact angle and bilayer area, which are often manipulated, in order to modulate the behaviour of the bilayer and transmembrane proteins ^27^. An example includes mechanosensitive protein channels, whose activation relies on the tension across the phospholipid bilayer ^28–30^. This has been achieved using chemical means, such as the hydrolysis of lysolipids by phospholipase A_2_ or through physical actuation of the membranes ^30,31^. Another bilayer manipulation example includes the concentration minimization of proteins pores and channels in DIBs, by directly dragging/pulling the droplets using electrodes or pipettes ^27,28,32^. Others have induced liquid volume-assisted pressure changes within the DIB droplet-based compartments, therefore manipulating the droplet size and the bilayer area ^33^. Alternatively, DIB manipulation has been achieved via electrowetting methods ^24^, or through the incorporation of magnetic particles and exposure to magnetic fields ^34^. Electrowetting manipulation of DIBs can be limited by electroporation and bilayer rupture ^35^, while mechanical manipulation can be constrained by the contact and movement of invasive pipettes and electrodes, often causing failure of the DIBs.

In this work, we propose a simple chemical approach to alter hydrogel-encapsulated DIB networks, to directly modulate the properties of artificial cells and enable the construct’s dynamic response to environmental changes. This concept is demonstrated by constructing eDIBs and observing their interaction with water-soluble lysophosphatidylcholine (LPC) for prolonged periods. These lysolipids are single-tailed phospholipids, which alter the surface tension of lipid monolayers and induce pressure changes along the phospholipid leaflet, as evidenced by artificial cell studies ^36–38^. We find that at high concentrations (10-fold higher than the critical micellar concentration), LPC ruptures the artificial membranes and promotes rapid release of the enclosed aqueous content. At low, sub-micellar concentrations the droplet network endures physical changes, with significant alterations to the contact angle and bilayer area. Lysolipids were able to provide a facile and indirect contact approach for determining the fate of enclosed DIBs in aqueous environments.

## Results and Discussion

### High-order, gel-encapsulated DIBs using monolithic 3D-printed microfluidic devices

Three, in series, droplet-forming microfluidic junctions facilitated the formation of encapsulated DIBs in hydrogel capsules (Fig. 1a). For planar microfluidic devices, the wettability is vital for successful and stable emulsion formation, which is usually achieved through channel surface modification, including plasma treatment and coatings^39^. Here, triple emulsion capsules were produced using a 3D-printed microfluidic device made from Nylon and Cyclin Olefin Copolymer (COC) without any surface treatment or other device post-processing. Nylon and COC polymers are known for their hydrophilic and hydrophobic surface property, respectively ^40,41^. The surface water contact angle measurements of 3D-printed Nylon and COC substrates exhibited water contact angles of 46 ° and 78°, respectively (Fig. 1b, i.-iii.). The print settings of each material (*SI Appendix, Table S1*) were kept consistent between all 3D-printed samples and microfluidic devices, as they can affect the final water contact angle of the substrate ^42^. The Nylon and COC microfluidic components fused well together with no indication of leaking when the humidity was controlled while printing. It should be noted that Nylon fibres and films have been previously used in digital and paper microfluidics as superamphiphobic and anti-corrosive substrates ^43–45^, however Nylon is not widely used in droplet-microfluidics or 3D-printed microfluidics, possibly due to its hygroscopic properties ^46^. Here, the 3D-printed Nylon microfluidic component offered a novel and facile method of producing high-order emulsions. In fact, Nylon filament is more suitable for dual-material 3D-printed microfluidic devices, since previously reported PVA devices were soluble in water, which limited the duration of the microfluidic experiments ^19^. Earlier established eDIB models have been generated using glass capillary/3D-printed hybrid microfluidic devices ^17^, or using double emulsion 3D-printed devices ^19^.

**Fig. 1.**
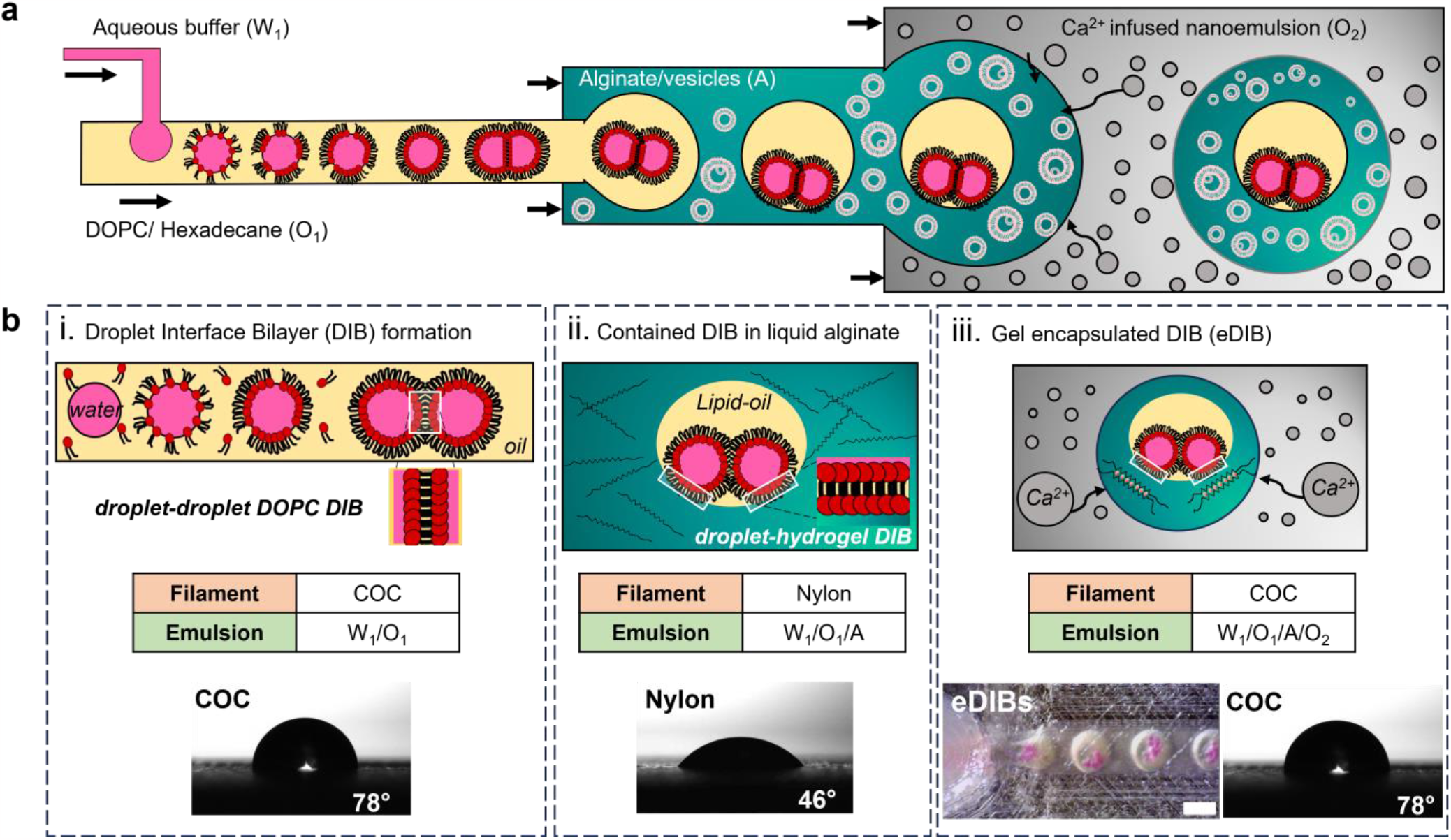
Monolithic 3D-printed microfluidic device generates triple emulsion capsules of encapsulated droplet interface bilayers (eDIBs). **a** Schematic of the triple emulsion microfluidic flow and production of eDIBs. The water phase (W_1_) is broken into droplets by the lipid-containing hexadecane oil (O_1_), which is then engulfed by a vesicle-containing alginate solution (A). The eDIBs are formed at the final 3^rd^ junction and gelled downstream by the Ca^2+^-infused nanoemulsion (O_2_). **b** Schematics of the stepwise generation of eDIBs from **a** including filament type, contact angle and emulsion order. i. Water-in-oil (W_1_/O_1_) emulsion formed by the 1^st^ hydrophobic (COC, 78°) droplet-forming junction. When the DOPC lipid monolayer-coated droplets come in contact, they form a DOPC droplet interface bilayer (DIB). ii. A close look at a water-in-oil-in-alginate (W_1_/O_1_/A) emulsion formed at the 2^nd^ hydrophilic (Nylon, 46 °) droplet-forming junction. The DIB is contained by an alginate phase with DPPC vesicles (vesicles are not shown). Where an inner aqueous droplet contacts the alginate, another DIB is formed defined as a droplet-hydrogel DIB. iii. Finally, the eDIB is formed at the 3^rd^ droplet-forming junction (COC, 78 °). The DIB contained by the alginate is engulfed by the Ca^2+^-infused nanoemulsion (W_1_/O_1_/A/O_2_), where the on-chip gelation starts (scale bar: 1 mm).

The final microfluidic devices consisted of three droplet-forming junctions. The 1^st^ and 3^rd^ junctions were made of COC filament and the 2^nd^ junction was made of Nylon filament. Initially, a water-in-oil (W_1_/O_1_) emulsion was formed at the 1^st^ droplet-forming junction, which advantageously exploited the COC filament’s hydrophobic properties. In the oil (hexadecane), DOPC phospholipids were dissolved and resulted in the formation of a lipid monolayer around individual water droplets, which when in contact with each other, formed DIBs (Fig. 1b, i.). Subsequently, the W_1_/O_1_ was inserted at the 2^nd^ droplet-forming junction made of Nylon filament (hydrophilic), and was broken by a continuous aqueous alginate phase. Therefore, multiple water droplets in lipid-containing oil (DIBs) were encapsulated in the liquid alginate, forming a water-in-oil-in-alginate (W_1_/O_1_/A) emulsion. At the site where an inner droplet comes in contact with the alginate, a droplet-hydrogel DIB is formed (Fig. 1b, ii.).

The lack of synthetic surfactants within the alginate resulted in the failure of the complex emulsion W_1_/O_1_/A (data not shown). Instead, of adding a surfactant into the alginate solution, we explored the addition of multilamellar DPPC vesicles, as surface tension-lowering agents ^47,48^. This hindered the coalescence between miscible phases. Both DOPC and DPPC phospholipids have been used towards the construction of artificial cell membranes (e.g. liposomes), hence either DOPC or DPPC could be used in the alginate phase, however, only DPPC vesicles were studied here. Finally, the W_1_/O_1_/A was encapsulated by a divalent-infused nanoemulsion, for further emulsification (W_1_/O_1_/A/O_2_) and simultaneous on-chip gelation (Fig. 1b, iii.). The final constructs are referred to as eDIBs, as they are hydrogel-based constructs encapsulating DIBs and can be stored in an aqueous environment. To our knowledge, this is the first report of fabricating monolithic, 3D-printed microfluidic devices that can generate multi-compartment triple emulsions microgels, without performing any device post-fabrication treatment or processing.

Free-standing eDIB capsules were produced with varying numbers of inner droplets. By controlling the flow rates of the inner aqueous buffer and the DOPC containing hexadecane phase we produced eDIBs with either an average diameter of 90 μm ± 1 μm (Fig. 2a) or 190 μm ± 3 μm (Fig. 2b). For reducing the diameter of the inner droplets, the aqueous phase flow rate of the inner droplets was decreased to 0.1 mL/hour, and the lipid-containing oil was increased to 0.5 mL/hour. eDIBs with smaller inner aqueous droplet diameters (ᴓ < 100 μm) have been shown to be notably more robust after centrifugation (*SI Appendix, Fig. S1*).

**Fig. 2.**
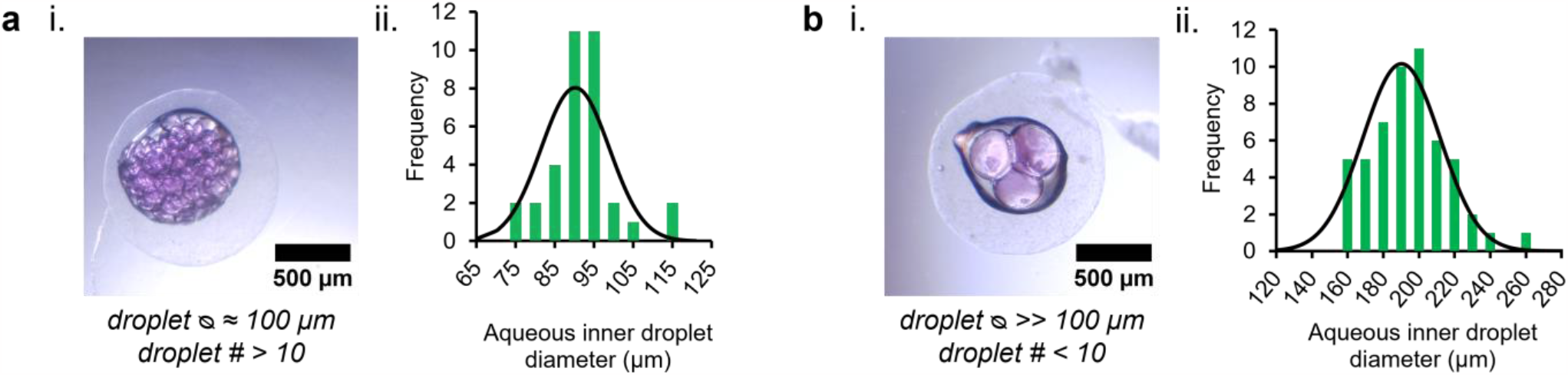
Gelled eDIB capsules with varying inner droplet diameter and number. **a** i. eDIB capsule containing many droplets (# > 10) of small diameter (ᴓ < 100 μm). ii. Inner droplet diameter distribution plot of the eDIB in i. (n=35). **b** i. eDIB capsule containing a small number of droplets (# < 10) of large diameter (ᴓ > 100 μm). ii. Inner droplet diameter distribution plot of the eDIB in i. (n=53). The gelled eDIBs shown were produced using the same microfluidic circuit and different flow rate combinations. Flow rates of **a** and **b** were 0.1 (W_1_): 0.5 (O_1_): 5 (A): 8 (O_2_) mL/hour and 0.2 (W_1_): 0.2 (O_1_): 5 (A): 8 (O_2_) mL/hour, respectively.

It should be noted that often with 3D FFF printed micro-scale components, variabilities may be introduced on the microfluidic channel dimensions (*SI Appendix, Table S2 and Fig. S2*), due to different environmental conditions and calibration inaccuracies. Because of these variabilities, eDIBs were formed at multiple phase flow rate combinations across experiments (*SI Appendix, Table S3*). For subsequent experiments the flow rates were manipulated accordingly, in order to enclose droplets with large diameter (ᴓ > 100 μm) and a small droplet number (typically less than 10), which would permit good visualization of the droplet arrangement and DIBs. eDIBs that survive the initial 2-3 hours of production can be stored for a month in an aqueous buffer with osmolarity that matches their internal droplets.

### Lysolipid-induced droplet release from eDIBs

Egg lysophosphatidylcholine (LPC) is a water-soluble, cone-shaped, single-tailed phospholipid with a headgroup larger than the tail, which tends to form micellar lipid structures with positive curvature ^49^. LPC has been used to alter the membrane pressure and activate mechanosensitive channels in DIB systems ^30^, increase the permeability of cell membranes for drug uptake studies ^50,51^, and facilitate protein pore insertion into bilayers ^52^. Here, the LPC lysolipid was introduced to the physiological aqueous environment surrounding the eDIB capsules and diffused passively to the phospholipid DIB between the inner aqueous droplets and the hydrogel shell (droplet-hydrogel DIB).

Prior to imaging, the eDIBs were immobilised at the bottom of a 96-well plate using 1 % w/v agarose, and this was followed by the addition of LPC in buffer at the final concentration of interest (Fig. 3a). The amphiphilic lysolipids diffused to the droplet-hydrogel DIB and at high concentrations (e.g. 100 μM) the inner droplets completely leaked into the surrounding medium, leaving an empty oil core (Fig. 3b). This was further analysed in terms of the fluorescent signal drop over time, across a population of eDIB capsules exposed to various LPC concentrations (1 μM – 1000 μM). The droplet release profile for each concentration over a period of 14 hours is shown in Fig. 3c. After approximately 3 hours of incubation at 37 °C and constant humidity, the intensity of 0 μM and 1 μM LPC treated eDIBs stabilised with negligible reduction. This reduction of the fluorescent signal was attributed to possible photobleaching and out-of-focus imaging, caused by the moving platform. In addition, the inner droplets of eDIB capsules treated with 10 μM and 100 μM LPC were subject to major instabilities after approximately 2-3 hours of the introduction of LPC. After the initial 3 hours, the 10 μM LPC-treated eDIBs were able to maintain their stability for longer time periods compared to 100 μM LPC-treated eDIBs. The logarithm of the intensity revealed exponential decay over time with fluctuations at concentrations of 10 μM and higher, whilst it also uncovered the bursting events at concentrations of 1000 μM.

**Fig. 3.**
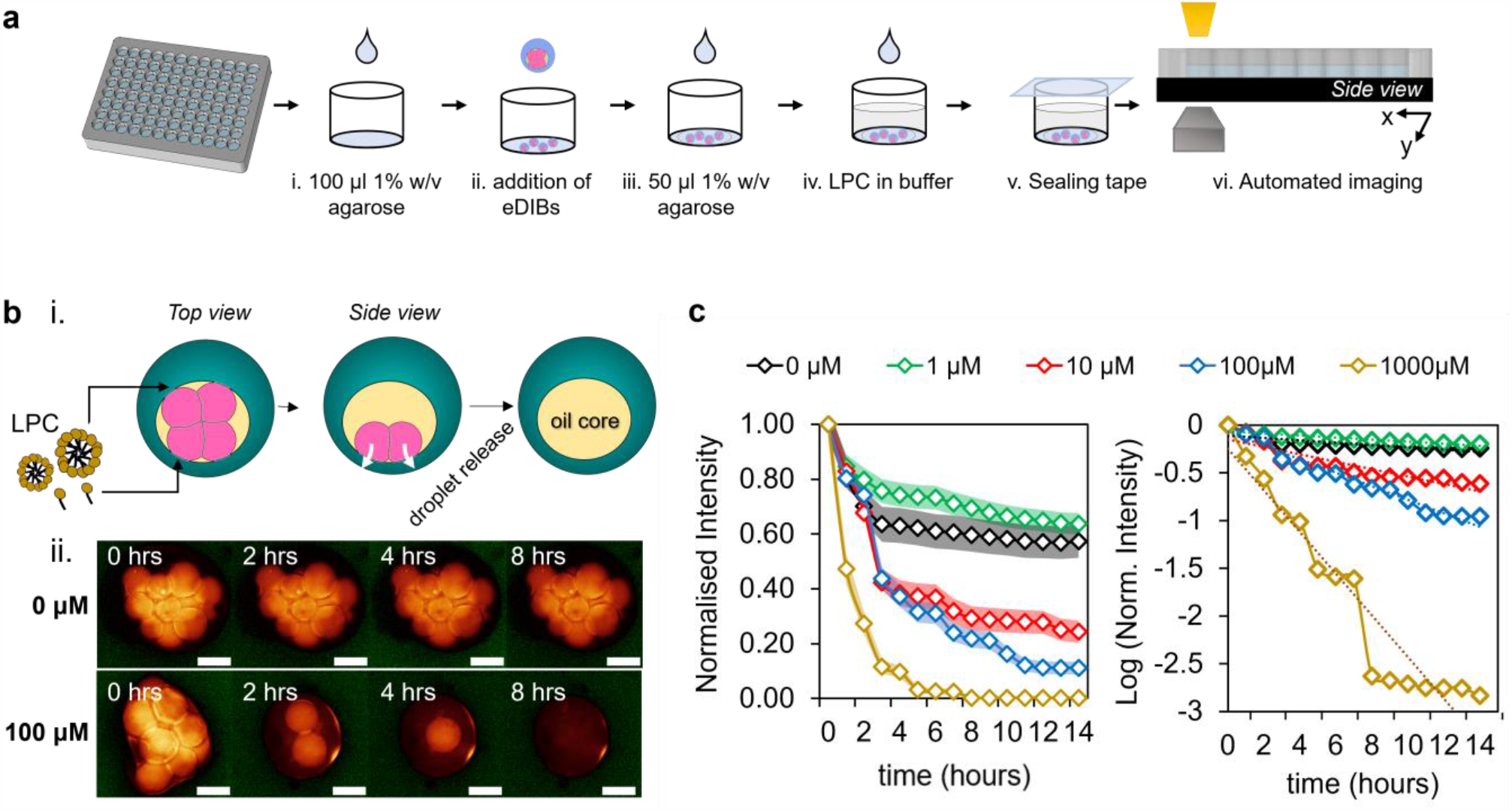
The effect of externally added LPC lysolipids on the release of inner aqueous droplet from eDIB capsules. **a** Stepwise schematic of the LPC treatment execution on eDIB capsules. First, a thin layer of 1 % w/v agarose was added to the bottom of the well, followed by the addition of eDIB capsules and then another thin layer of agarose. This facilitated the immobilization of eDIBs at the bottom of the plate during the treatment and imaging with the EVOS automated platform. The temperature of the imaging platform was kept at 37 °C and the humidity was controlled by a well plate sealing tape. **b** i. Top view and side view schematic of the eDIB capsules, showing the external addition of monomeric and micellar LPC. During the incubation of the eDIBs with concentrated LPC micelles, the lysolipids interact with DIBs formed between the hydrogel and inner aqueous droplets (droplet-hydrogel DIB) and subsequently, the droplets get released into the hydrogel. ii. Time-lapse of the aqueous fluorescent (sulforhodamine B) inner droplets, showing the rapid release from eDIBs treated with micellar LPC concentrations (100 μM). Scale bar: 200 μm. **c** Fluorescent signal of the eDIB inner droplets incubated with different concentrations of LPC (fluorescent decrease assay). The intensity reduction for the untreated eDIB capsules (0 μM) is attributed to artefacts of the automated imaging platform and photobleaching. The sample population per concentration for the intensity analysis was as follows: n= 11 (0 μM), n=15 (1 μM), n=19 (10 μM), n=17 (100 μM), n=16 (1000 μM). LHS: Normalised intensity versus time. The shaded regions for each line plot correspond to the standard error of mean (±SEM). RHS: The normalised intensity replotted in the logarithmic (log) scale over time. Besides this exponential decay, there are three consistent fluctuations at concentrations 10 μM, 100 μM and 1000 μM showing a small delay with decreasing concentration. These fluctuations begin during a secondary process and finally level out.

The phosphatidylcholine composition used in this study was dominated by approximately 69 % of 16:0 Lyso PC (information provided by manufacturer), leading to the assumption that the critical micelle concentration (CMC) is close to that of 16:0 Lyso PC (CMC_LPC_). The CMC value is a variable of temperature, pH and salt ^53,54^, and the exact CMC_LPC_ was not measured in this study. However, previous literature reported that the CMC_LPC_ value of 16:0 Lyso PC ranges between 4 μM and 8.3 μM, at temperatures spanning from 4 °C to 49 °C ^55,56^. Therefore, only the 10 μM concentration introduced to the eDIBs in this study, was considered as a concentration closest to previously reported CMC_LPC_.

Either individual LPC lipid molecules, monomers (<CMC_LPC_), or micelles (>CMC_LPC_) were delivered to the droplet-hydrogel DIB and interacted with the first outer leaflet of the bilayer. This will alter the curvature of the membrane, leading to an asymmetric pressure distribution along the bilayer ^57^. High micellar concentrations of LPC can lead to the rupture of phospholipid membranes, as a consequence of the translocation of crowded lysolipids to the second inner leaflet of the bilayer, or due to lysolipid-induced perturbations ^51,58,59^. Similarly, here the droplets treated with equal to or greater than 100 μM LPC were subject to rapid droplet bursting, due to the failure of the droplet-hydrogel DIB membrane. In comparison to lower concentrations, this active release was attributed to the concentrated LPC micelles delivered to the targeted site (droplet-hydrogel DIB) and promptly induced membrane asymmetry. Supplementary fluorescence increase assays showed that 10 μM treated eDIBs underwent a major droplet-hydrogel DIB failure at a later timepoint (∼7 hours), compared to higher concentrations which caused instant membrane failure (*SI Appendix, Fig. S3*).

For lysolipids to diffuse and act on the droplet-hydrogel DIB, the monomers and micelles need to diffuse from the aqueous solution and then through the alginate shell. Lysolipids can interact and fuse with the DPPC lipid vesicles embedded in the hydrogel alginate shell, leading to the possible reduction of the lysolipid fraction delivered to the droplet-hydrogel DIB. An underestimated lysolipid concentration can influence the rate of impact on the eDIB constructs, which explains why the effects occur in the order of hours. Furthermore, the micellar size highly depends upon the concentration, where 7-50 μM LPC form micelles of 34 Å radius, whereas this micellar radius doubles at concentrations exceeding 50 μM ^60^. Consequently, concentrations equal to or higher than 100 μM deliver large micelles, which contribute to the possible transient pore formation at the bilayer, thus the droplet-hydrogel DIB instantly fails and droplet release into the hydrogel occurs ^36,61^.

### The effect of sub-micellar LPC concentrations on droplet displacement and arrangement

Lipid monolayer-coated aqueous droplets in the form of water-in-oil emulsion are governed by the interfacial tension. Bilayer and DIB formation is facilitated by Van-der-Waals forces, as the adhesive monolayer-coated droplets come in contact ^62^. Due to the excess of lipids in the hexadecane oil phase, the monolayers of a DIB can expand and contract. There are temporary fluctuations of the disjointing pressure during DIB formation at the various interfaces, but attractive and repulsive forces work towards the equilibrium of the eDIB system ^63–65^.

Significant fluctuations begin when the lysolipids are externally introduced, as illustrated in Fig. 4a. The equilibrium is destabilized, due to the arrangement of the introduced lysolipids into the existing phospholipid bilayer and the subsequent changes in the pressure distribution ^66^. The reorganization of the phospholipids within the first encountered lipid monolayer of the droplet-hydrogel bilayer, results in a change in the surface tension and subsequent lateral expansion, as lysolipids are being fed into the monolayer (Fig. 4a, ii.). Meanwhile, this forces the excess DOPC phospholipids in the hexadecane oil to compensate from the internal side of the droplet-hydrogel bilayer, towards the LPC-induced monolayer expansion. Together, both leaflets of the bilayer endure tensional changes, which causes the adhesive forces at the droplet-hydrogel bilayer to shift and the whole bilayer to expand along the interface.

**Fig. 4.**
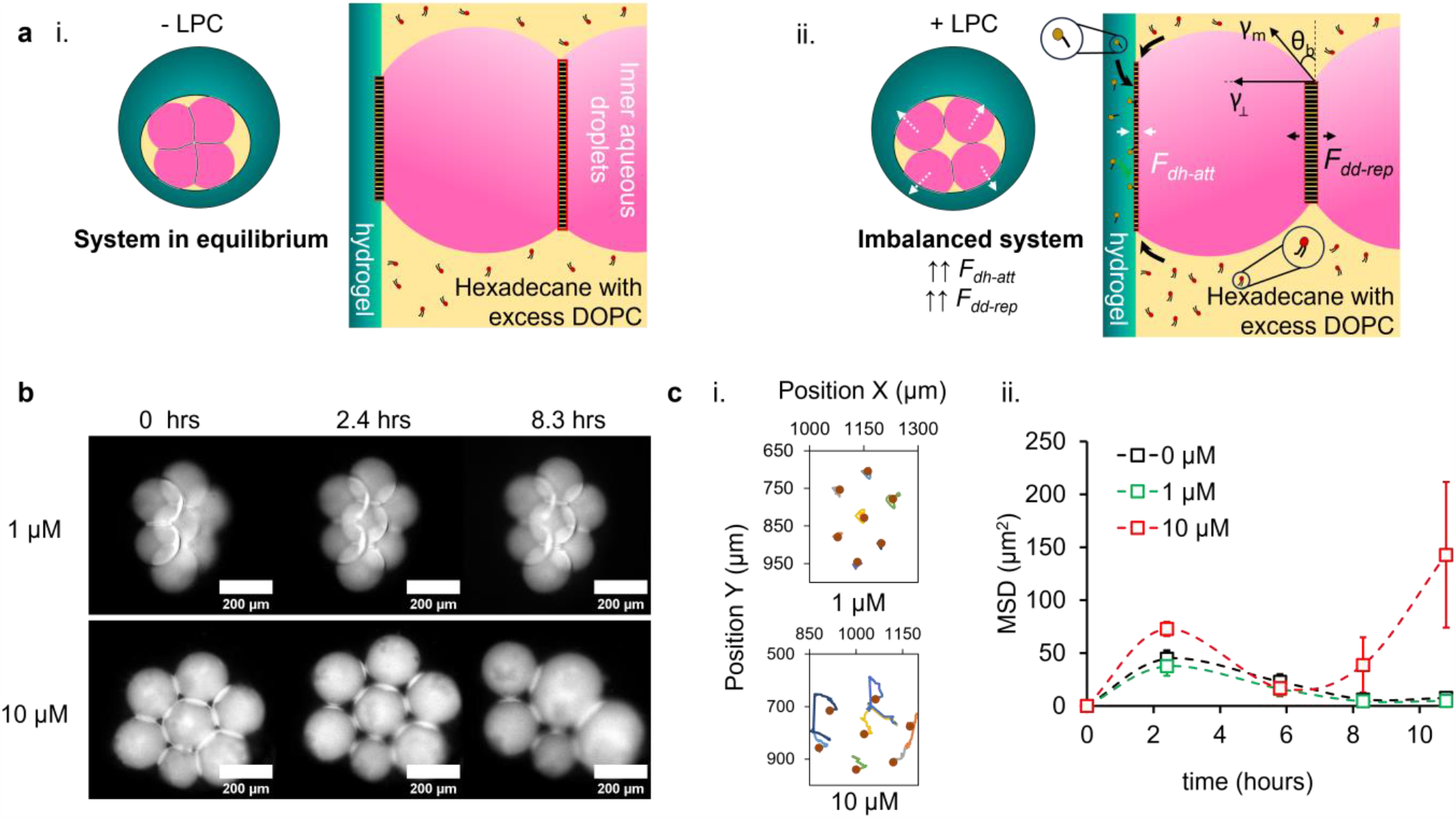
Inner droplet dynamics and re-arrangement under the influence of sub-micellar LPC lysolipid concentrations. **a** Schematic diagram of eDIBs and key bilayer interfaces before (-LPC) and after (+ LPC) the addition of lysolipids. i. The eDIB system and bilayer interfaces are at equilibrium, as attractive and repulsive forces balance each other. ii. The introduced lysolipids take the eDIB out of equilibrium, as the LPC and DOPC contribute to the lateral expansion of the droplet-hydrogel DIB, by inserting from the external and internal side of the bilayer, respectively. Consequently, the attractive forces at the droplet-hydrogel DIB 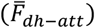 rise, which in turn drive the repulsive forces at the droplet-droplet DIB 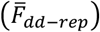. These forces, 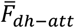 and 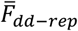, lead to the thinning and thickening of the droplet-hydrogel DIB and droplet-droplet DIB, respectively. The contact angle (θ_b_) between the droplets is influenced by the increasing repulsive forces and the characteristic surface tension, *γ_⊥_*. **b** Time lapse of the inner aqueous droplets of eDIBs treated with 1 μM and 10 μM LPC, showing significant pulling and subsequent merging of droplets treated with 10 μM LPC. **c** Plots of the, i. X and Y position of the inner droplets and, ii. the mean square displacement (MSD) of 0 μM, 1 μM and 10 μM LPC treated eDIBs measured over 11 hours, revealing that 1 μM treated droplets travelled similar to the untreated construct, while there was significant travel by 10 μM treated droplets. The dots in i. show the location of the individual droplets at t=0. Error bars in ii. correspond to the standard error of mean (±SEM).

In the presence of lysolipids, we classify the dominating forces of the eDIB model as the attractive forces at the droplet-hydrogel DIB 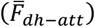, and repulsive forces at the droplet-droplet DIB 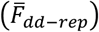. As the droplet-hydrogel bilayer laterally expands in the presence of LPC, the 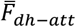 dominate over any other attractive forces and the bilayer becomes thinner (Fig. 4a). This leads to the pulling of the droplets towards the hydrogel shell, due to imbalanced forces. Here, we describe the pulling effect as the retraction of the droplets away from the centre of the middle oil core. At this stage, 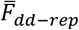 are dominating and the thickness of the droplet-droplet bilayer increases. Above a characteristic critical bilayer thickness the aqueous droplets will separate, while below a critical bilayer thickness, coalescence occurs ^65^. Otherwise, an equilibrium might be reached when the forces balance each other, and a characteristic equilibrium bilayer thickness is achieved.

The above molecular dynamic changes after the addition of LPC promote the rearrangement and displacement of the inner droplets and DIBs. The rate and the degree of destabilization effects depended on the concentration of LPC introduced. Droplet pulling was more explicit in eDIBs treated with 10 μM LPC, as shown in Fig. 4b. eDIBs treated with 1 μM LPC were overall less disturbed with mean square displacement similar to the control (0 μM), while the displacement of the droplets exposed to 10 μM LPC was more apparent (Fig. 4c). After approximately 8 hours of incubation with the lysolipids, the pulling effect led to droplet merging for eDIBs treated with 10 μM LPC. In fact, during the study period and at this concentration of LPC, it was observed that the inner droplets would initially merge between them, and not with the hydrogel shell. This was due to the enhanced stability of DIBs formed on hydrogel semi-flat substrates ^67^, compared to droplet-droplet DIBs. Once the first merging occurred, a cascade of merging continued where small droplets merged with larger droplets (the product of merging), due to the higher Laplace pressure inside smaller droplets ^67^. The delayed droplet shifting and displacement in the presence of 10 μM LPC (Fig. 4c, ii) were attributed to the slower build-up of lysolipid concentration at the droplet-hydrogel DIBs ^68^.

### The effect of sub-micellar LPC concentrations on DIB bilayer area and contact angle

High bilayer tension and strong adhesion forces at the droplet-hydrogel DIB contributed to the increased bilayer area and contact angles ^23^. The bilayer area and contact angle were not captured at the droplet-hydrogel DIB, due to imaging limitations, and were only measured between the aqueous droplet-droplet DIBs.

The three-dimensional micro-architecture of the eDIB capsules benefited the measurements of circular bilayer areas, which reflect the shape of the droplets, throughout incubation as shown in Fig. 5a. This allowed the quantification of circular bilayer areas of DIBs between adjacent droplets, which helps assess the bilayer stability and behaviour ^16^. The bilayer area of 1 μM treated eDIBs shows delayed effects induced by LPC and subsequent return to equilibrium, as evident by the bilayer area plateau (Fig. 5a, ii). Moreover, the bilayer area of vertical droplet-droplet DIBs inside eDIB capsules was calculated on the assumption that the droplets on either side of the bilayer were of equal diameter (*SI Appendix, Fig. S4)*. Fig. 5b shows the average bilayer area of 1 μM and 10 μM treated DIBs throughout the incubation period. Whilst negligible bilayer area reduction was observed with 1 μM LPC, the 10 μM LPC caused a significant reduction within the initial 3.5 hours, followed by droplet merging (bilayer area increase) and then once again, bilayer area reduction.

**Fig. 5.**
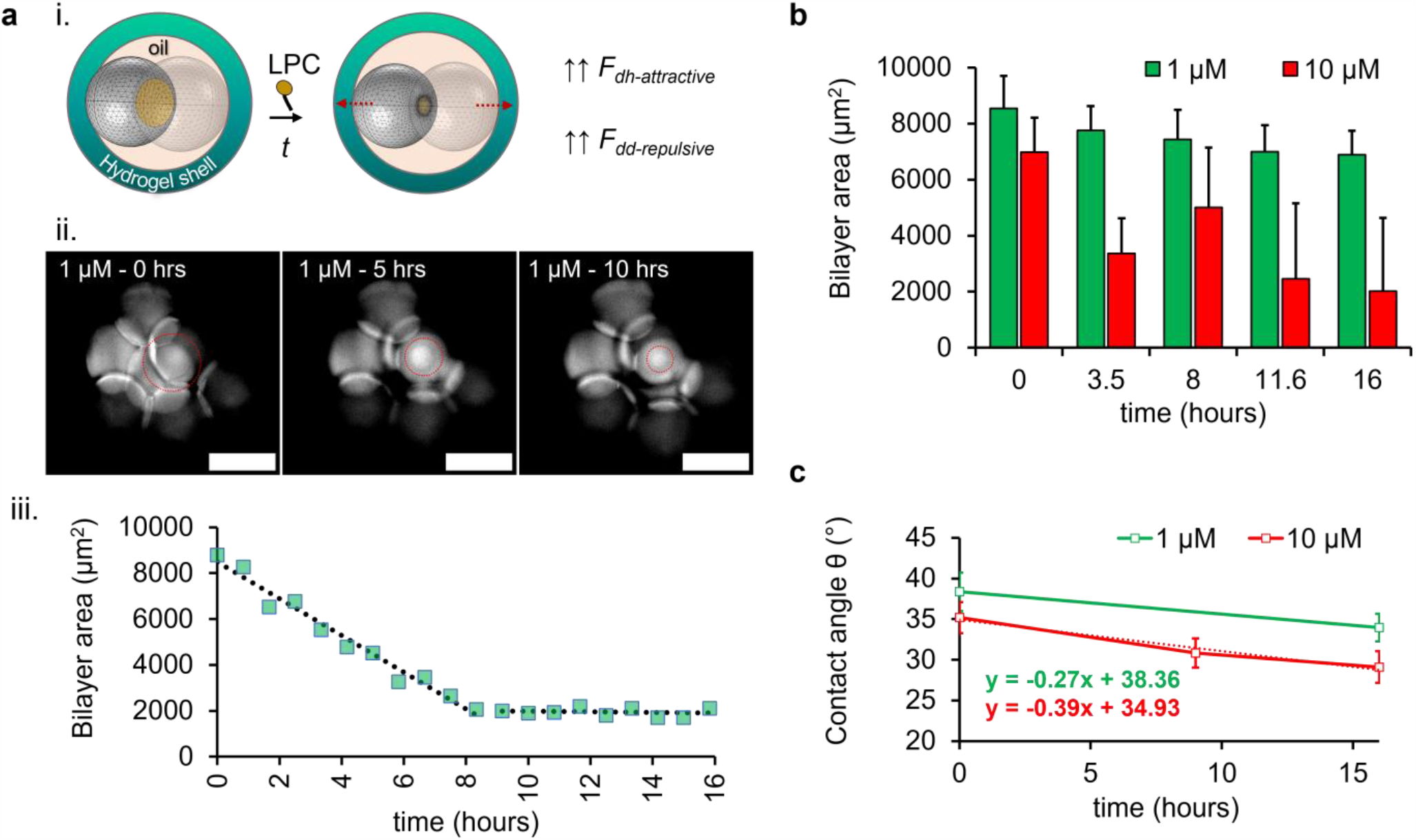
LPC lysolipid impact on the bilayer area and contact angle of eDIBs. **a** i. A schematic of an eDIB capsule with two inner droplets and a formed DIB (yellow circular droplet contact area), before and after the addition of LPC. The DIB area is reduced during incubation with LPC, as the adhesive forces of the droplet-hydrogel bilayer and repulsive forces at the droplet-droplet bilayer begin to dominate (*↑↑* F_dh-attractive_, *↑↑* F_dd-repulsive_), ii. Time-lapse of fluorescent droplets encapsulated within an eDIB capsule treated with 1 μM LPC, showing the reduction of the bilayer area as indicated by the red dotted circle. To reveal the bilayer between the contacting droplets, the brightness and contrasts of the image were adjusted. Scale bar: 200 μm. iii. The measured circular bilayer area from ii. is plotted over time as a scatter plot, whilst the dotted curve shows the linear decrease in the first 8 hours after 1 μM LPC addition; this is followed by a transition to a constant bilayer area (equilibrium reached) until the end of the study. **b** Average DIB bilayer area over time across a population of eDIBs treated with 1 μM (n=11, N=4) and 10 μM (n= 12, N=5) LPC. The DIB bilayer area of 10 μM treated constructs displays a drop at 3.5 hours and then an increase at approximately 8 hours, which indicates first the pulling of the droplets and subsequent merging, respectively. After that, the bilayer area follows a reduction and begins to equilibrate. A minimal and subtle decrease was observed in the bilayer area throughout the study in 1 μM treated eDIBs. The number of measured vertical bilayers for 10 μM treated DIBs was initially n= 12 (N=5), and this dropped to n=4 (N=5) by the final timepoint, due to droplet merging. **c** Line graph of the average DIB contact angle as a function of time for 1 μM (n =55, N= 6) and 10 μM (n= 47, N=9) treated eDIB capsules. An additional timepoint at approximately 9 hours was plotted, which corresponds to the initial merging of droplets treated with 10 μM (best fit for 10 μM treated eDIBs shown by the dotted line). The line plots are accompanied by linear equations, which reveal the initial average DIB contact angle (38 ° for 1 μM and 35 ° for 10 μM). The population number of the measured contact angles for 10 μM was n=47 (N=9), and this dropped to n=22 (N=9) by the final timepoint, due to droplet merging. The number of eDIBs is noted by N, whilst the sample population of the measurable characteristic (bilayer area or contact angle) is noted by n.

The contact angle of DIBs typically depends on the droplet diameter, lipids and oil composition, as they can affect the surface tension and consequently the droplet-droplet adhesion ^21^. The number of droplets enclosed within a volume forming DIBs can also affect the droplet-droplet contact angle ^23^. In most DIB models, the contact angle of DIBs is manipulated prior to the DIB formation by varying the lipid and oil composition (*SI Appendix, Fig. S5)*. In this study, the contact angle among the inner compartments can be manipulated post-fabrication through the incubation of the eDIBs with sub-micellar LPC concentrations. This is displayed in Fig. 5c., where the mean contact angle between eDIB droplets was measured before the LPC started to affect the droplet-droplet DIBs to a measurable extent, and at the end of the incubation. In addition to the endpoint contact angle measurements, the contact angle was measured at approximately 9 hours for eDIBs only treated with 10 μM LPC, representing an average timepoint after the first droplet coalescence.

Bilayer peeling between two droplets of contact angle *θ_b_* and monolayer surface tension was previously attributed to the exceeding of the critical adhesive bilayer force per unit length, by a quantity *γ_⊥_* which drives the droplet-droplet DIB separation, *γ_⊥_ = γ_m_* sin *θ_b_* ^25^ (Fig. 4a, ii.). Here, the peeling or pulling at the droplet-droplet DIB is indirectly driven by the dominating monolayer surface tension, attractive forces and droplet shape deformation at the droplet-hydrogel DIB.

Overall, we demonstrated the successful encapsulation of droplet interface bilayer membranes into self-supported hydrogel capsules (eDIBs) using COC/Nylon, 3D-printed microfluidic devices. This was benefited by utilizing lipid vesicles as interfacial tension-altering particles, which hindered the mixing between miscible phases. In addition, a method was established for inducing and monitoring the release of the inner aqueous compartments from these free-standing, complex emulsion capsules using micellar lysolipid concentrations. Sub-micellar concentrations, on the other hand, induced more refined effects, including 3D reorganisation and changes in the bilayer area and contact angle.

Advantages for employing microfluidics in DIB model construction include the high production yield, and control over the size and structural order, whilst various features can be introduced, such as phospholipid bilayer asymmetry. The incorporation of phospholipid DPPC vesicles within the alginate phase can contribute towards the formation of asymmetric DOPC/DPPC bilayers, following partial lipid-in and lipid-out DIB formation. Although, in this study, we considered a lipid-out symmetric DOPC bilayer constructed at the bilayer interface between droplets, and between any droplet and the hydrogel.

An earlier active-release study on hydrogel eDIBs demonstrated the synergy between pore-forming peptides, but no control over the organization or DIB adhesion was reported ^69^. Furthermore, when cholesterol molecules insert between the phospholipids of a bilayer, they create a condensed monolayer with restricted motion between the acyl chains of the phospholipids ^65^. Similar to cholesterol molecules, lysolipids at non-pore-forming concentrations insert between phospholipid molecules and increase the surface tension of the phospholipid monolayer (for LPC, tension will be higher between polar headgroups) and the energy of adhesion of the bilayer.

For the duration of the lysolipid LPC treatment, we hypothesized that the LPC molecules introduced to the eDIB system were unable to encounter the droplet-droplet DIB, directly. Therefore, the lysolipids only affected the droplet-hydrogel bilayer, where strong adhesion forces pull the droplets and attenuate the droplet-droplet DIB area. These findings present an approach for in-situ and automated organization, as well as the manipulation of the bilayer area and contact angle of encapsulated droplet-droplet DIBs. It should be noted that, the duration of phospholipid bilayer exposure to lysolipids can enhance the lipid molecular transfer to the opposite leaflet ^70^ and hence, the degree of impact.

Research in artificial cells and protein reconstitutions would benefit from the non-invasive modulation of artificial cellular membranes. Simply by introducing lysolipids we can modulate the spatial organisation and physicochemical characteristics of encapsulated droplet interface bilayers. These findings pave the way for non-invasive transmembrane protein density control studies, as well as establishing communications between the internal and external environment of artificial cell chassis. Complex artificial membrane models such as eDIBs aided by droplet-microfluidics offer the benefit of encapsulating and interfacing two or more reagent-carrying compartments. Droplet microfluidic technology provides a versatile tool for developing such increasingly sophisticated droplets structures for artificial cellular models to study biomolecular interactions and precision engineering of encapsulated bioinspired membranes.

## Materials and Methods

### Materials

COC was purchased from Creamelt (Grade 8007, TOPAS) and transparent Nylon was purchased from Ultimaker. Sulforhodamine B and calcein were purchased from Thermofisher, UK. The calcein and sulforhodamine were dissolved in 0.05 M HEPES, 0.15 M KCl in deionised water (buffer) or Phosphate Buffered Saline, PBS (pH 7.4, 1 X, Gibco, UK). Alginic acid sodium salt from brown algae, hexadecane, silicone oil AR20, mineral oil, calcium chloride, HEPES, potassium chloride, 1,2-di-oleoyl-sn-glycero-3-phosphocholine (DOPC), 1,2-dipalmitoyl-sn-glycero-3-phosphocholine (DPPC), Egg lysophosphatidylcholine (LPC), chloroform and SPAN 80 were purchased from Merck. The average fatty acids in the egg lysophosphatidylcholine mixture according to the manufacturer was 69 % 16:0, 24.6 % 18:0, 3.4 % 18:1, 1.4 % 16:1, 0.3 % 14:0, 0.3 % 18:2 and 1 % unknown.

### 3D-printed microfluidic device fabrication and operation

The microfluidic device was designed using COMSOL Multiphysics (versions 5.6) and fabricated using the Ultimaker S5 Pro Bundle with cyclic olefin copolymer (Creamelt) and Nylon (Ultimaker). The device was sliced using the CURA software with the assigned print settings summarized in *SI Appendix*. All devices after printing were stored with silica gel sachets. Each liquid phase was delivered to the microfluidic device using SGE gas-tight glass syringes loaded onto positive displacement syringe pumps (KD Scientific). The SGE syringes were connected directly to the 3D printed microfluidic inlets using PTFE tubing (O.D. ᴓ = 1.58 mm, I.D. ᴓ = 0.80 mm). Further details regarding the microfluidic device, channel dimensions and flow operation can be found in *SI Appendix*.

### Production of Water-in-Oil-in-Water-in-Oil eDIB capsules (W_1_/O_1_/A/O_2_)

All reagents were purchased from Merck, unless otherwise stated. The inner water phase (W_2_) consisted of a buffer solution of 0.05 M HEPES, 0.15 M potassium chloride, 200 μM of sulforhodamine B (SulfB) or 70 mM calcein. The middle oil phase (O_1_) consisted of 12.5 mg/mL 1,2-di-oleoyl-sn-glycero-3-phosphocholine (DOPC) in hexadecane. DOPC was first dispersed in hexadecane following the thin film lipid hydration method. Briefly, the DOPC powder was dissolved in chloroform and evaporated using a gentle nitrogen stream until a thin film of lipids was formed. The DOPC film was subject to a vacuum for at least 30 minutes to evaporate any residual chloroform and then released under nitrogen gas. The shell phase (A) consisted of 1 % w/v alginate and 0.5 mg/mL 1,2-dipalmitoyl-sn-glycero-3-phosphocholine (DPPC) vesicles in buffer. The DPPC vesicle solution was prepared using the thin film lipid hydration method, following vacuum overnight. The DPPC film was dispersed in the buffer solution, vortexed for 30 seconds and sonicated in a water bath at 55 °C for 15 minutes. The eDIB capsules’ oil carrier phase (O_2_) consisted of a Ca^2+^ - infused mineral oil emulsion, which facilitated the gelation of the alginate shell. This carrier phase was prepared by mixing an aqueous solution of 1 g/mL CaCl_2_ and mineral oil at 1:9 volume ratio, with 1.2 % SPAN 80 surfactant. The mixture was stirred for at least 10 minutes using a magnetic stirrer and plate, creating a Ca^2+^ - infused nanoemulsion. During experiments, the outlet orifice was slightly submerged in 0.2 M CaCl_2_.

The microfluidic setup and execution here, aimed at the formation of approximately 1 mm diameter eDIBs, with large water droplet compartments (> 100 μm) segregated by artificial lipid membranes (i.e., DIBs).

### LPC treatment of eDIBs

eDIB capsules were immobilized with 1 % w/v low temperature melting agarose in wells of a 96-well plate. LPC in buffer was prepared and used appropriately, in order for each well to have final LPC concentration of 1, 10, 100 and 1000 μM. The droplet release was evaluated by monitoring the decrease in the fluorescence of sulfB (200 μM) from the droplets of individual eDIBs or the fluorescence increase in the wells with eDIBs encapsulating quenched calcein (70 mM). Details related to the LPC fluorescence increase assay can be found in the *SI Appendix*.

### Optical and Fluorescence Microscopy of eDIBs

eDIBs during on-chip emulsification were imaged using Dino Lite edge USB microscope. eDIBs post-production and during the LPC treatment were imaged using EVOS M7000 Imaging System. Imaging associated with the lysolipid treatment was carried out at 37 °C, where the well plate containing the eDIB capsules was sealed with a tape to prevent evaporation.

### Bilayer area and DIB contact angle measurements

The bilayer area was measured in three different ways depending on the bilayer orientation and sphericity of the droplets forming the DIB. See *SI Appendix* for full bilayer area calculation. Due to the ability of the inner droplets to maintain their three-dimensionality, the contact angle was simply calculated by measuring the angle between two adjacent inner aqueous droplets using the angle drawing tools on ImageJ. Before measuring the angle, the contrast of the image was adjusted accordingly, in order to remove any noise around the region of interest. The bilayer area and contact angle of eDIBs produced using 4 mg/mL DOPC in 10 % silicone oil were also measured as reference and comparison to conventionally produced eDIBs (12.5 mg/mL DOPC, 100 % hexadecane).

### Fluorescence and image analysis

The droplet release in the fluorescence decrease assay was evaluated by monitoring the fluorescence decrease from the aqueous sulfB droplets of individual eDIBs (measured the area of the fluorescent DIBs inside the whole construct.). The droplet release in the fluorescence increase assay was evaluated by monitoring the fluorescence increase of the wells with the eDIBs carrying droplets of quenched calcein (70 mM). Image handling and fluorescence analysis were carried out using ImageJ software. The integrated fluorescent intensity was measured at the timepoint of interest, with the ROI minimized to the area of the fluorescent droplets. The intensity plots show the intensity normalised to the intensity extracted from the control (0 μM LPC) fluorescent droplets in eDIBs. The position and displacement of the droplets was recorded using manual tracking tools within ImageJ. The eDIB samples were monitored for over 10 hours and the position of the droplets was recorded every 5 minutes.

## Acknowledgments

This work was partially supported and funded by the European Horizon 2020 project ACDC (Artificial Cells with Distributed Cores) under project award number 824060.

