## Supplementary Information for "Manipulation of encapsulated artificial phospholipid membranes using sub-micellar lysolipid concentrations"

**Supporting Information**

**3D-printing settings.** All microfluidic devices were designed using COMSOL Multiphysics (versions 5.6) and fabricated using the Ultimaker S5 Pro Bundle (Ultimaker S5 installed with the air manager and material station). Each design was exported from COMSOL Multiphysics software as an .STL file and imported into CURA slicing software. In CURA, the print settings were assigned as summarized in Table S1, to generate a G-CODE file for the 3D printer. All devices after printing were stored in silica gel sachets. A thin layer of PVA glue was applied to the glass platform of the 3D printer, before device fabrication. The Nylon was kept inside the material station of the Ultimaker S5 Pro Bundle, in order to minimize moisture absorption.

**Operation of microfluidic devices and flow.** Each liquid phase was delivered to the microfluidic device using SGE gas-tight glass syringes loaded onto positive displacement syringe pumps (KD Scientific). The SGE syringes were connected directly to the 3D-printed microfluidic inlets using PTFE tubing (O.D. ᴓ = 1.58 mm, I.D. ᴓ = 0.8 mm). A small amount of UV resin was applied to each inlet to seal the fluidic connection and was cured with a UV torch (365 nm). The microfluidic flows from day-to-day experiments were imaged using a Dino-Lite Edge USB microscope, unless otherwise stated. The microfluidic devices that served eDIB production were based on the same CAD design. However, the 3D FFF printing of microfluidic devices often suffered from small dimensional inaccuracies, as shown by the comparisons between the 3D-printed dimensions and CAD dimensions (Table S2 and Fig. S2). For this reason, it was not feasible to use the exact same flow rate per device. eDIBs were produced by microfluidic devices operated at a range of flow rates, shown in Table S3.

Small and tightly packed DIBs were associated with increased stability ^1^, and this was shown by centrifugation experiments (Fig. S1), although, the microfluidic setup and execution here, aimed at the formation of approximately 1 mm diameter eDIBs, with large water droplet compartments (> 100 μm).

**COC and Nylon contact angle measurements.** Contact angle measurements for COC (Creamelt, Grade 8007, TOPAS) and Nylon 3D-printed samples were collected according to BS EN 828:2013. OneAttension Theta Lite optical tensiometer was used as the contact angle measuring system. The samples were printed as blocks with dimensions 3 cm x 3 cm x 0.1 cm, using the Ultimaker S5 Pro Buddle and the same print settings as the microfluidic devices (Table S3). The samples were printed on a glass platform, after a thin layer of PVA glue was applied. Each block sample was cleaned after fabrication with DI water and 70% ethanol to remove residual PVA glue and dust. Each sample was positioned on the OneAttension platform and a 2 μl water drop was placed on the surface of the sample. Measurements of the contact angle were taken by the camera over 10 s, starting as soon as the 2 μl water drop was deposited on the sample. The surface of four (n=4) 3D-printed block samples of COC and Nylon were tested for the contact angle. The reported average contact angles for COC and Nylon were calculated based on measurements over 1 s (every 0.05 s), immediately after the water droplet touches the surface.

**Lipid-particle assisted eDIB formation.** During preliminary experiments, the lack of surfactants led to merging between miscible phases upon the entry into the subsequent junction or channel. This occurred between the inner water droplets (WP) and the alginate phase (AP) or, between the mid-oil phase (OP) and the outer oil carrier phase (OC). These observations arise from the merging of miscible phases within the microfluidic chip in the absence of surfactants. To combat the coalescence during on-chip microfluidic formation of eDIBs, others have added CaCO_3_ particles in the alginate phase ^2^. Here, we minimize the effect of coalescence by incorporating DPPC lipid-vesicles into the alginate phase (AP). DPPC lipid vesicles reduced the surface tension of alginate ^3,4^, and acted as small particles that supported lipid bilayer formation ^5,6^. Additionally, lipid vesicles can act as emulsion stabilizing agents, by developing a barrier that mitigates merging effects between miscible phases. This stabilizing property of lipid particles inspired the addition of lipid vesicles to the alginate phase, in order to facilitate the generation of eDIBs.

**LPC treatment fluorescence increase assay (quenched dye).** Self-quenched calcein at a concentration of 70 mM was encapsulated in the inner aqueous droplets of eDIBs. The droplet release assay was the same for all LPC treatments. Five LPC concentrations were tested on eDIBs encapsulating the quenched dye (0 μM, 1 μM, 10 μM, 100 μM, 300 μM). The LPC treatment on eDIBs encapsulating sulforhodamine B, were subject to 1000 μM LPC, instead of 300 μM. The eDIBs exposed to the two highest concentrations (100 μM and 300 μM) underwent faster release than the concentration closer to the CMC (10 μM) (Fig. S3).

**Bilayer area measurements.** Gravity is neglected and the droplets on either side of the bilayer are assumed to have equal diameter. Z-stack images of the eDIBs used for the bilayer area measurements were obtained with a step size of 17.2 μm.

*Area of vertical bilayers formed by spherical droplets (Fig. S4(a-b))*

The area of vertical bilayers formed between spherical droplets was calculated based on the length of the vertical bilayer $\left( a_{bil} \right)$. This vertical length may be subject to errors due to the focal plane (< 17.2 μm). Due to the sphericity of the droplets, the bilayer is also assumed to have a spherical shape, hence the bilayer area is calculated as:

$$A_{bil}=\frac{\pi{a_{bil}}^{2}}{4}$$

Equation S1

*Area of vertical bilayers formed by ellipsoid droplets (Fig. S4(c-d))*

The area of vertical bilayers of DIBs formed between ellipsoid droplets was calculated by introducing the eccentricity parameter $\left( \varepsilon\right)$, which describes the deviation of an ellipsoid from a circle. The eccentricity parameter of a droplet $\varepsilon_{drop}$, considers the minor $\left( b_{drop} \right)$ and major semi-axis $\left( a_{drop} \right)$ of the ellipsoid:

$$\varepsilon_{drop}= \sqrt{\left( 1-\left( \frac{b_{drop}}{a_{drop}} \right)^{2} \right)}$$

Equation S2

The bilayer interface between ellipsoid droplets can undergo the same deformation as the droplets ^7^, hence:

$$\varepsilon_{drop}= \varepsilon_{bil}= \sqrt{\left( 1-\left( \frac{b_{bil}}{a_{bil}} \right)^{2} \right)}$$

Equation S3

Where, $\varepsilon_{bil}$ is the eccentricity of the bilayer interface. Rearranging and substituting Equation S3 into the area of an ellipsoid, the bilayer area can be expressed in terms of the major semi-axis of the vertical bilayer $\left( a_{bil} \right)$ and $\varepsilon_{bil}$:

$$A_{bil}=\frac{\pi{a_{bil}}^{2}\sqrt{1-\varepsilon_{bil}^{2}}}{4}$$

Equation S4

*Area of horizontal bilayers formed by spherical droplets*

eDIBs that were oriented in a way where the horizontal bilayer was visible to the objective, the bilayer area of horizontal bilayers was simply measured by the circular tool from ImageJ, which technically employes Equation S1.

**Table S1.** COC and Nylon print settings assigned within CURA software for Ultimaker S5 3D printer. These setting were used for all experiments and microfluidic devices.

| **Print Setting** | **COC*** | **Nylon*** |
| --- | --- | --- |
| **Speed** | 25 mm/s | 20 mm/s |
| **Infill** | 100 % (0.69 mm) | 100 % (0.69 mm) |
| **Initial Layer Height** | 0.18 mm | 0.18 mm |
| **Layer height** | 0.06 mm | 0.06 mm |
| **Line width** | 0.22 mm | 0.23 mm |
| **Wall line width** | 0.22 mm | 0.23 mm |
| **Material Flow** | 100 % | 100 % |
| **Fan Speed** | 100 % | 10 % |
| **Printing temperature** | 255 °C | 245 °C |
| **Build plate temperature** | 85 °C | 85 °C |

*Brim Tower for Dual Printing was enabled

**Table S2.** Comparison of the channel dimensions between CAD design dimensions and the fabricated 3D-printed channels. Each letter corresponds to the same channel dimension of the CAD designs and the optically measured microfluidic channels from Fig.S2. The channels were designed as cylinders (**ᴓ**), unless marked by the symbol of , implying that the particular channel was designed as a rectangle.

|  | **b** | **d** | **e ᴓ** | **g ᴓ** | **i ᴓ** | **k ᴓ** |
| --- | --- | --- | --- | --- | --- | --- |
| **Design dimensions** | 0.36 mm | 0.50 mm | 0.50 mm | 1 mm | 1.40 mm | 1 mm |
| **3D-printed device dimension** | 0.24 mm | 0.30 mm | 0.45 mm | 0.86 mm | 1.36 mm | 0.95 mm |
| **% difference** | 12 % | 20 % | 5 % | 14 % | 4 % | 5 % |

**Table S3.** Fluid phases and flow rates used for eDIB production of this study. One example of flow rate combination is 0.1 (WP), 0.2 (OP), 3 (AP), 8 (OC) mL/hour.

| **Liquid Phase** | **Description** | **Flow rates (mL/hour)** |
| --- | --- | --- |
| **Aqueous inner droplets with dye (WP)** | Buffer or PBS with calcein or sulfB | 0.1-0.5 |
| **Lipid-containing oil (OP)** | 12.5 mg/mL phospholipids in hexadecane | 0.2-0.8 |
| **Hydrogel shell (AP)** | 1.5 % alginate, 0.5 mg/mL DPPC | 3-6 |
| **Nanoemulsion (OC)** | 1:9 Ca^2+^-infused mineral oil | 5-12 |

**Fig. S1. Centrifuged eDIB capsules with fluorescent droplets of diameter less than 100 μm.**

Dark field images taken at **a** 0 hours and at **b** 48 hours post centrifugation. The small droplets sit at the bottom of the DOPC/hexadecane oil core, forming tightly packed networks with no indication of leakage into the hydrogel. As the centrifugal force increases, the top surface of the DIB networks becomes flatter. The eDIBs were centrifuged in 1 mL of mineral oil.

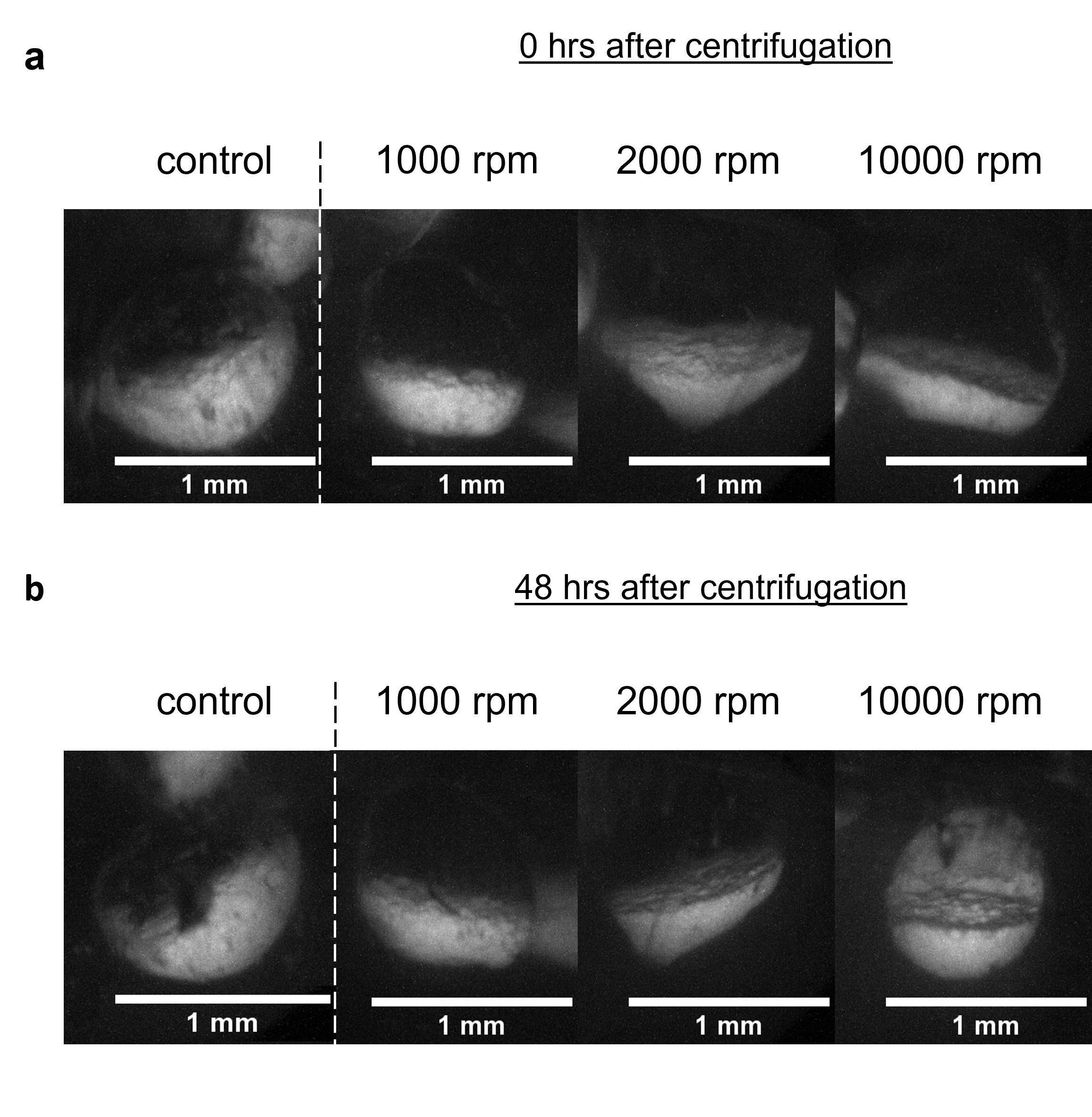

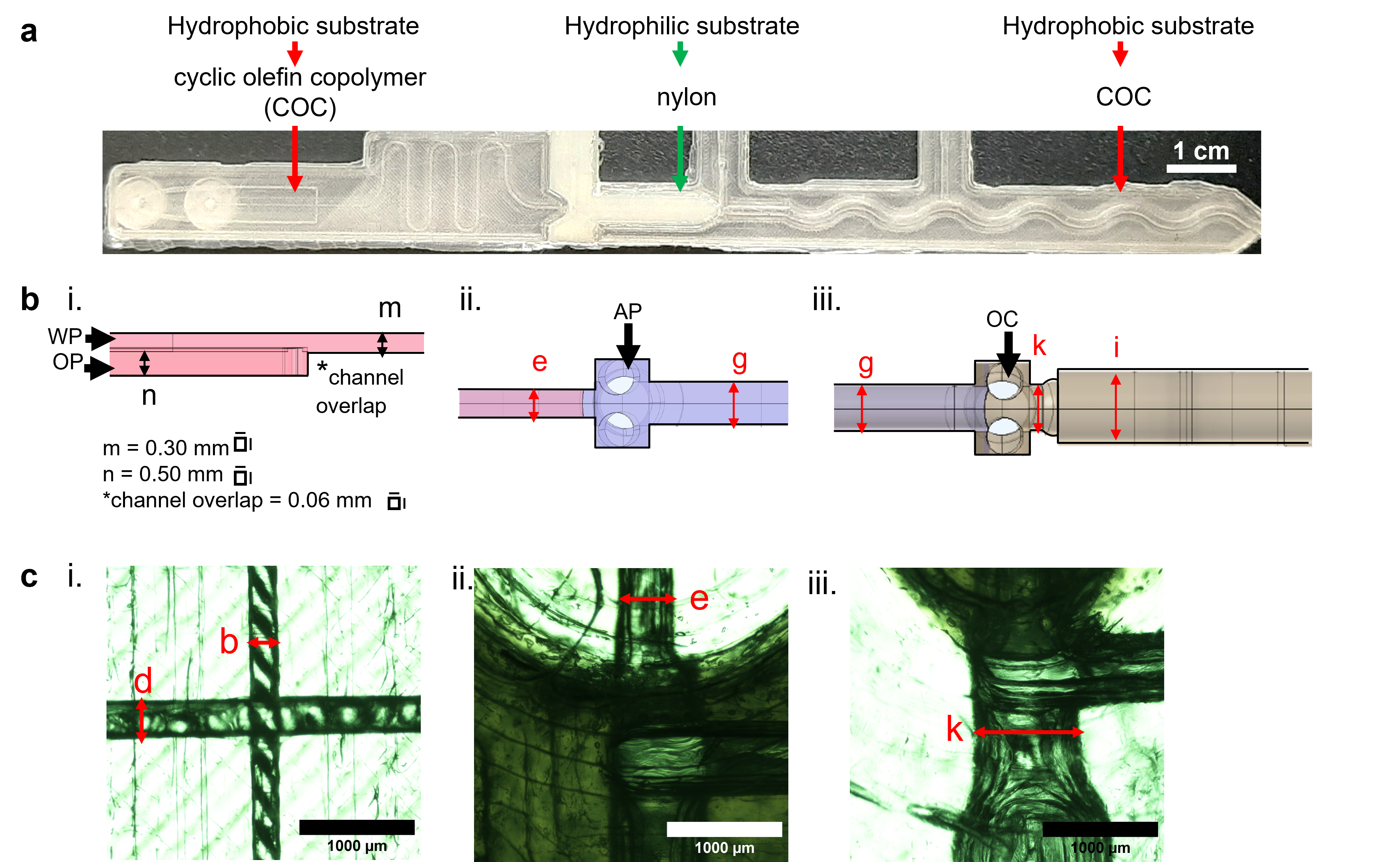

**Fig. S2. Microfluidic device characterisation and dimensions.**

**a** An image of the 3D-printed microfluidic device utilised for the formation of eDIBs. Triple emulsion formation is achieved by fabricating droplet-forming junctions with materials of opposing wettability, e.g., COC and Nylon, which are considered hydrophobic and hydrophilic respectively. **b** Side view of the CAD design of the microfluidic device. i. Two-plane COC 1^st^ droplet-forming junction for the production of the inner aqueous droplets or DIBs (W/O), ii. 2^nd^ droplet-forming junction made of Nylon for W/O/W emulsion and iii. 3^rd^ droplet-forming junction for engulfing the whole eDIB by the calcium-infused oil carrier (OC) phase. **c** Top view images of the 3D-printed droplet-forming junctions obtained using widefield microscopy. i. 1^st^ droplet-forming junction, where b and d are the inner water phase (WP) and lipid-oil phase (OP), respectively. Then follows the ii. 2^nd^ droplet-forming junction and iii. 3^rd^ droplet-forming junction. The Nylon components of the device appear darker compared to COC, due to Nylon’s reduced transparency. The dimensions of the channels noted by the red letters can be found in Table S3. The channels were designed as cylinders (**ᴓ**), unless marked by the symbol of , implying that the particular channel was designed as a rectangle.

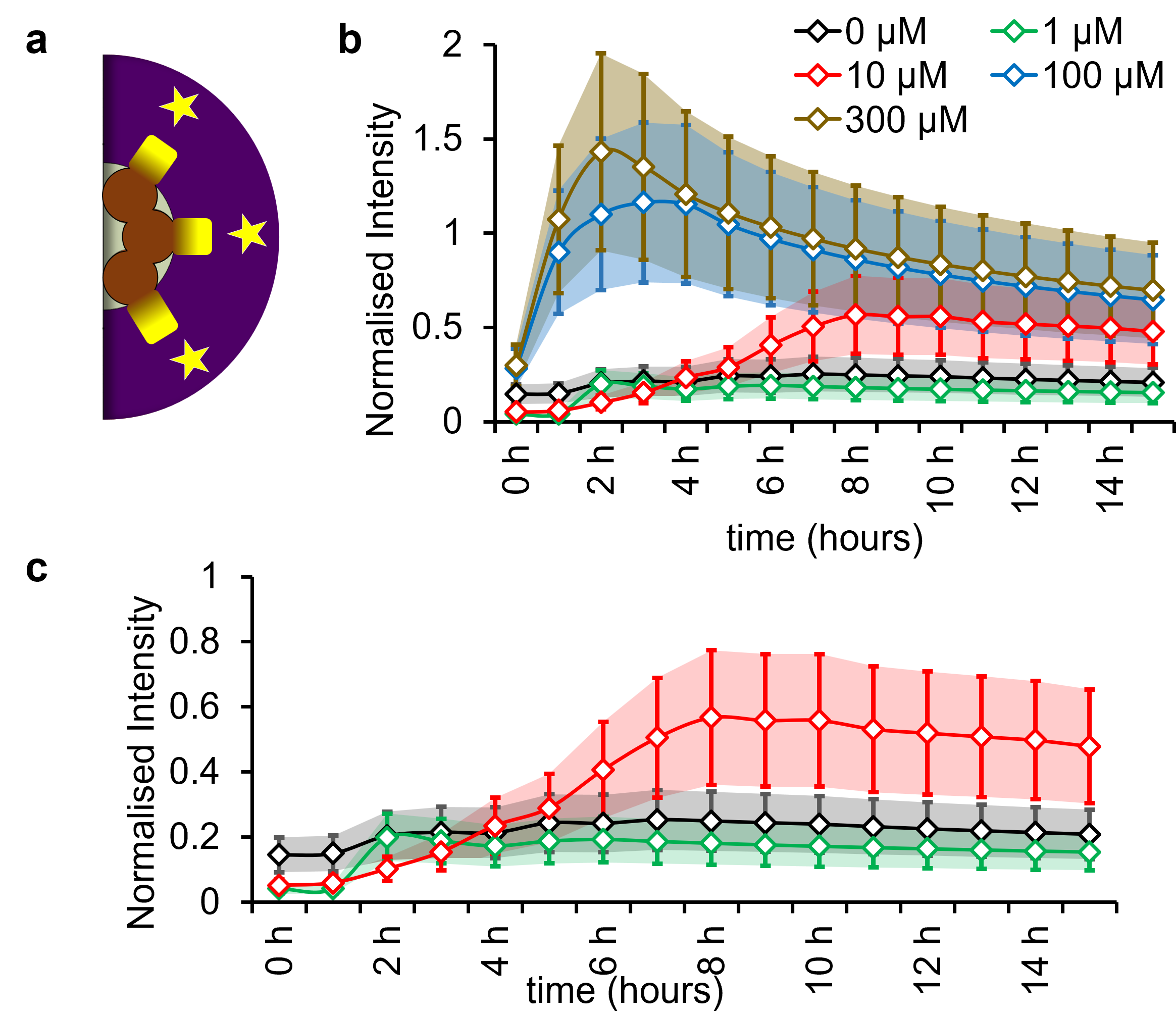
**Fig. S3. Fluorescence increase assay of eDIBs encapsulating quenched calcein dye, treated with LPC.**

**a** Schematic of an eDIB, showing that when the inner aqueous droplets are released under the influence of LPC, the surrounding becomes fluorescent (yellow stars). **b** Fluorescent intensity profile of the eDIBs incubated with different concentration of LPC. Self-quenched calcein at a concentration of 70 mM was encapsulated in the aqueous droplets of eDIBs. This plot shows rapid release by eDIBs treated with 100 μM and 300 μM LPC. The sample population per concentration for the intensity analysis is as follows: n= 10 (0 μM), n=5 (1 μM), n=3 (10 μM), n=3 (100 μM), n=3 (1000 μM). **c** Closer look on the quenched calcein eDIBs treated with 0 μM, 1 μM and 10 μM LPC. Strong fluorescence was detected at approximately 8 hours after the addition of 10 μM LPC, which is attributed to the initial droplet-hydrogel membrane failure.

**Fig. S4. DIB schematics between spherical and elliptical droplets.**

The noted dimensions where used to calculate the bilayer area of DIB formed by (**a**-**b**) spherical droplets and droplets that deviated from the spherical shape, due to the silicone oil (**c**-**d**) (α_bil_=bilayer diameter, b_drop_=minor semi-axis of the droplet, α_drop_=major semi-axis of the droplet).

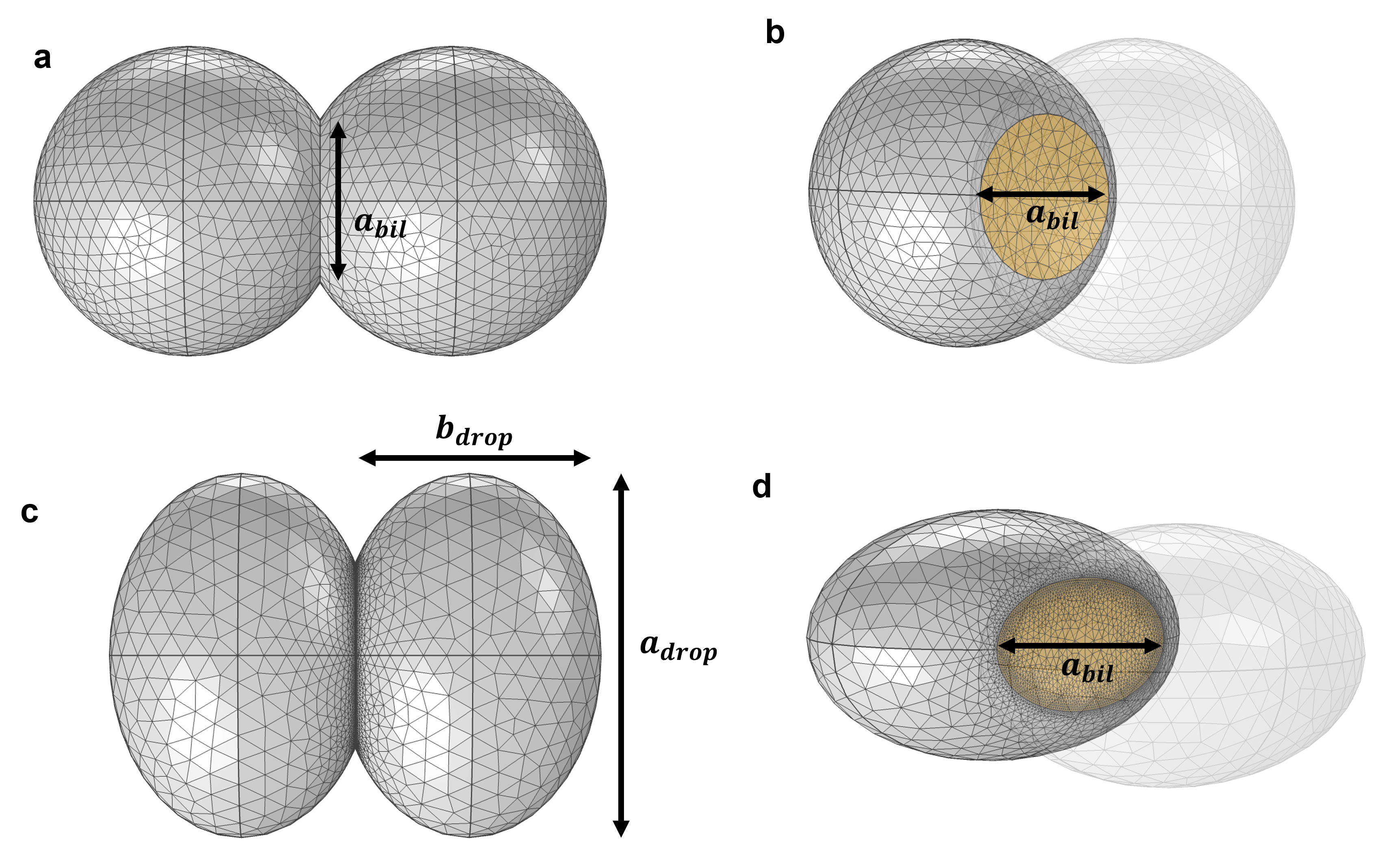

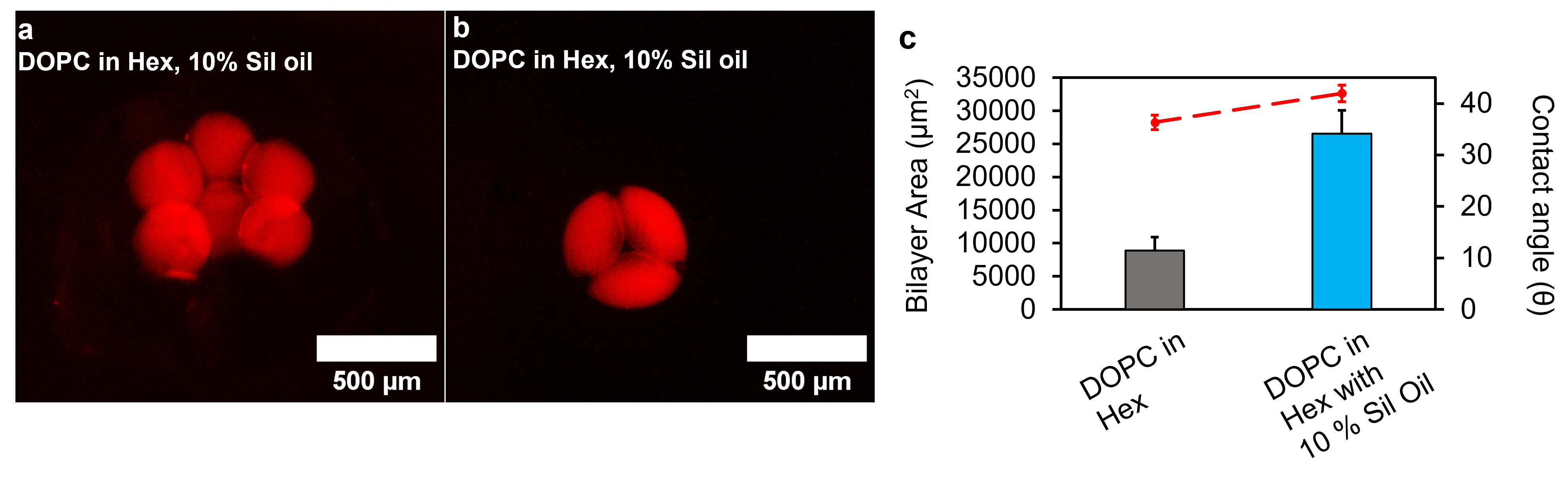
**Fig. S5. eDIBs of two lipid-containing oil compositions.**

**a** 12.5 mg/mL DOPC in 100 % hexadecane, **b** 4mg/mL DOPC in hexadecane with 10 % silicone oil. **c** Bilayer and contact angle bar and line plot, respectively, for eDIBs produced with either DOPC in hexadecane only, or DOPC in hexadecane with 10 % silicone oil. The average contact angle of eDIBs formed with 10 % Silicone oil (4mg/mL DOPC in hexadecane) was approximately 41.9 ° ± 1.4 °, while conventional eDIBs obtained a contact angle of approximately 36.6 ° ± 0.6 °. As the surface tension of the oil surrounding the droplets, i.e. DIBs, decreases in the presence of surface tension reducing oils (e.g. silicone oil), the droplets become less spherical, compared to pure hexadecane. The droplet’s shape deviation from a sphere subsequently affects the contact angle between droplets, leading to an approximate contact angle increase of 15 %, compared to pure hexadecane eDIBs. It should be noted that in other non-encapsulated DIB models, the contact angle was observed to be linearly proportional to the fraction of the silicone oil ^8^.
